# Precise quantification of behavioral individuality from 80 million decisions across 183,000 flies

**DOI:** 10.1101/2021.12.15.472856

**Authors:** Benjamin de Bivort, Sean Buchanan, Kyobi Skutt-Kakaria, Erika Gajda, Chelsea O’Leary, Pablo Reimers, Jamilla Akhund-Zade, Rebecca Senft, Ryan Maloney, Sandra Ho, Zach Werkhoven, Matthew A-Y Smith

**Affiliations:** Center for Brain Science & Dept. of Organismic and Evolutionary Biology, Harvard University, Cambridge, MA, USA; Dept. of Biology and Biological Engineering, California Institute of Technology, Pasadena, CA, USA; Department of Neurobiology, Harvard Medical School, Boston, MA, USA; Karius, Redwood City, CA, USA; Imaging Platform, Broad Institute of MIT and Harvard, Cambridge MA, USA; Department of Entomology, University of Wisconsin-Madison, Madison WI, USA

**Keywords:** handedness, fluctuating asymmetry, variability, high-throughput behavior, automation, ethology

## Abstract

Individual animals behave differently from each other. This variability is a component of personality and arises even when genetics and environment are held constant. Discovering the biological mechanisms underlying behavioral variability depends on efficiently measuring individual behavioral bias, a requirement that is facilitated by automated, high-throughput experiments. We compiled a large data set of individual locomotor behavior measures, acquired from over 183,000 fruit flies walking in Y-shaped mazes. With this data set we first conducted a “computational ethology natural history” study to quantify the distribution of individual behavioral biases with unprecedented precision and examine correlations between behavioral measures with high power. We discovered a slight, but highly significant, left-bias in spontaneous locomotor decision-making. We then used the data to evaluate standing hypotheses about biological mechanisms affecting behavioral variability, specifically: the neuromodulator serotonin and its precursor transporter, heterogametic sex, and temperature. We found a variety of significant effects associated with each of these mechanisms that were behavior-dependent. This indicates that the relationship between biological mechanisms and behavioral variability may be highly context dependent. Going forward, automation of behavioral experiments will likely be essential in teasing out the complex causality of individuality.

## Introduction

Individual animals exhibit idiosyncratic behavior, even when their genetics and rearing environment are held constant. This variability is termed intragenotypic variability (Stamps et al., 2013) and likely arises in part due to stochastic effects during development (Vogt 2015; Honegger & de Bivort, 2018), which, in a quantitative genetic framework, are classified as microenvironmental plasticity (Morgante et al., 2015). Intragenotypic variability in animal behavior is likely a major component of animal personality, an ecologically and evolutionarily important dimension of variation (Freund et al., 2013; Bierbach et a., 2017). A single genotype giving rise to a broad distribution of random phenotypes may constitute an adaptive evolutionary strategy, termed “bet-hedging,” to increase the probability that for any fluctuation in the environment, some individuals will be fit, increasing the odds that a lineage never goes extinct (Hopper, 1999). While bet-hedging has strong theoretical foundations, in the context of animal behavior it has limited evidence (but see Kain et al., 2015 and Akhund-Zade et al., 2021). A challenge in studying bet-hedging is that behavioral variability is difficult to measure; larger sample sizes are needed to precisely estimate the variance of a trait, compared to the mean. This is largely because the former requires sampling phenotypes in the tail of a distribution, which are rare by definition.

Increasing behavioral assay throughput via automation is an effective way to attain the sample sizes needed to study variability. This can be achieved through miniaturization and parallelization of imaging platforms in a lab context (Kain et al., 2012; Pantoja et al., 2016; Churgin et al., 2017; Stern et al., 2017; Barlow et al., 2021). While the up-scaling of experiments is easiest with small, lab-adapted animals, such approaches do work with species beyond the common genetic models (Crall et al., 2015; Bierbach et al., 2017; Crall et al, 2018; Ulrich et al., 2018). Gains in data throughput can be achieved with the help of robots that automate animal handling (Alisch et al., 2018), move cameras between arenas (Alisch et al., 2018; Crall et al., 2018) or track a single animal over long periods of time (Johnson et al., 2020). Automation of analysis is also essential, and innovations in animal centroid tracking (Panadiero et al., 2021), body-part tracking using neural networks (Hausmann et al., 2021) and behavioral classification (Kabra et al., 2013; Berman et al., 2014; Todd et al., 2017) constitute a rich tool set for rapidly extracting behavioral measures from digital data sets.

High-throughput, automated behavioral assays have been used to investigate the variability of *Drosophila* behavior (Steymans et al., 2021; Werkhoven et al., 2021; Mueller et al., 2021; Mollá-Albaladejo & Juan Sánchez-Alcañiz 2021). The species’ deep genetic toolkit facilitates the study of proximate mechanisms controlling variability such as neurotransmitters (Kain et al., 2012; Honegger & Smith et al., 2019), neural circuits (Buchanan et al., 2015; Honegger & Smith et al., 2019; Linneweber et al., 2020; Skutt-Kakaria et al., 2019), genes (Kain et al., 2012; Ayroles et al., 2015; Wu et al., 2018), environmental variation (Akhund-Zade et al., 2019), and social effects (Alisch et al., 2018; Versace et al., 2019). Of these studies, the three that have assayed the greatest number of individuals (Buchanan et al., 2015, Ayroles et al., 2015 and Skutt-Kakaria, et al., 2019) all employed a common behavioral assay: spontaneous locomotion in Y-shaped mazes. As flies walk freely in these arenas, they make a left-vs-right choice every time they cross through the center of the maze. Individual flies make hundreds of such choices per hour. This yields a large data set per individual, which in combination with a high throughput of individuals, makes this assay particularly amenable to the study of variability. Beyond the number of left-right choices made and their average handedness, the Y-maze assay also produces behavioral measures related to the higher-organization of turn sequences and their timing (Ayroles et al., 2015).

Individual left-vs-right turning bias is correlated with clockwise-vs-counterclockwise bias in open arenas (Buchanan et al., 2015) indicating that the behavioral measures in this assay are not entirely geometry-dependent. Humans may exhibit a comparable form of locomotor bias in the curvature of their trajectories when trying to walk straight without visual feedback (Souman et al., 2009). The left-right symmetry of this assay evokes the phenomenon of fluctuating asymmetry, in which individual variation in the extent of morphological asymmetry is used as a measure of developmental stability (Van Valen, 1962; Debat et al., 2011). Indeed, both left-vs-right turn bias in Y-mazes and morphological traits examined for fluctuating asymmetry tend to have average values (typically close to left-right symmetry) that are robust across genotypes and selection (Pelabon et al., 2005; Ayroles et al., 2015).

Here, we took advantage of the high precision and throughput of the Y-maze assay to characterize the distribution of individual behaviors and their variability along different experimental axes. We collected nearly all the data from Y-maze experiments conducted by lab members since this assay was devised in 2010. In descriptive analyses, we characterized the distribution of individual Y-maze behavioral measures, and their correlations, with unprecedented precision. In hypothesis-driven analyses, we examined the effects on variability of manipulations of serotonergic signaling, the gene *white* (previously shown to affect phototactic variability; Kain et al., 2012), sex, and temperature. On the whole, these analyses reinforce the finding that genotype and the choice of behavioral measure itself have consistently large effects on measures of variability (Akhund-Zade et al., 2019), though some mechanistic manipulations can have large effects in a behavior-dependent fashion.

## Results

We collected experimental records from hundreds of experiments examining the Y-maze behavior of 183,496 individual flies (Figure 1). In total, these flies made 79.8M left-right choices. Four behavioral measures were recorded for each fly (Ayroles et al., 2015): turn bias (percent of turns to the right), number of turns, and turn switchiness. The latter is a measure of the degree to which flies alternate between left and right turns, normalized by their turn bias. A fly making exactly as many left (right) followed by right (left) turns as expected in a binomial model has a switchiness value of 1. Lower switchiness indicates fewer LR/RL turn sequences, and, conversely, longer streaks of L or R turns. The fourth measure, turn clumpiness, captures the non-uniformity of turn timing, i.e., the extent to which flies made choices in bursts. We changed the formula for the last measure midway through the data collection period (compare Buchanan et al., 2015 and Werkhoven et al., 2021), making this measure hard to compare across experiments; therefore we excluded it from further analysis. In addition to behavioral data, the record for each fly also included metadata about the experimental circumstances, including (Table 1): the fly’s genotype, experimental conditions, temperature during behavior, age of the fly, the experimenter who recorded the behavioral data, the ID# of the array of arenas (“tray”) in which it behaved, the ID# of the imaging box in which it behaved, the date, the number of arenas in its tray, the software used to record its behavior, the software used to produce its behavior measures, and its sex. The proportion of all flies for five of these metadata categories are shown in Figure 1B.

**Table 1.** Y-maze data set variables.

| data variable name | notes |
| --- | --- |
| flyID | number linking this fly's data to other digital records |
| handedness | turn bias behavioral measure |
| numTurns | number of turns behavioral measure |
| switchiness | turn switchiness behavioral measure |
| lev_handedness | Levene-transformed turn bias, for linear modeling of variability in turn bias |
| lev_numTurns | Levene-transformed number of turns, for linear modeling of variability in number of turns |
| lev_switchiness | Levene-transformed turn switchiness, for linear modeling of variability in turn switchiness |
| genotype | genotype of fly |
| expCond | experimental condition |
| expTemp | temperature during behavioral experiment (°C) |
| 5htpagar | flies reared on agar media supplemented with 10mM 5-HTP |
| 5htpagar25 | flies reared on agar media supplemented with 25mM 5-HTP |
| 5htpagar50 | flies reared on agar media supplemented with 50mM 5-HTP |
| 5htpnormal | flies reared on cornmeal-dextrose media supplemented with 10mM 5-HTP |
| 5htppotato10 | flies reared on potato media supplemented with 10mM 5-HTP |
| 5htppotato25 | flies reared on potato media supplemented with 25mM 5-HTP |
| 5htppotato50 | flies reared on potato media supplemented with 50mM 5-HTP |
| aMWnormal | flies reared on cornmeal-dextrose media supplemented with 10mM aMW |
| agar | flies reared on control agar media |
| amwagar | flies reared on control agar media supplemented with 15mg/mL ascorbic acid |
| amwpotato10 | flies reared on potato media supplemented with 10mM aMW |
| amwpotato20 | flies reared on potato media supplemented with 20mM aMW |
| amwpotato25 | flies reared on potato media supplemented with 25mM aMW |
| amwpotato50 | flies reared on potato media supplemented with 50mM aMW |
| ctrlaanormal | flies reared on control potato media |
| ctrlaapotato | flies reared on control potato media supplemented with 15mg/mL ascorbic acid |
| d10gal80heatshock | flies subjected to heat-shock at day 10 of development (Ayroles et al., 2015) |
| d14gal80heatshock | flies subjected to heat-shock at day 14 of development (Ayroles et al., 2015) |
| d1gal80heatshock | flies subjected to heat-shock at day 1 of development (Ayroles et al., 2015) |
| d3gal80heatshock | flies subjected to heat-shock at day 3 of development (Ayroles et al., 2015) |
| d4gal80heatshock | flies subjected to heat-shock at day 4 of development (Ayroles et al., 2015) |
| d5gal80heatshock | flies subjected to heat-shock at day 5 of development (Ayroles et al., 2015) |
| d6gal80heatshock | flies subjected to heat-shock at day 6 of development (Ayroles et al., 2015) |
| d7gal80heatshock | flies subjected to heat-shock at day 7 of development (Ayroles et al., 2015) |
| d8gal80heatshock | flies subjected to heat-shock at day 8 of development (Ayroles et al., 2015) |
| d9gal80heatshock | flies subjected to heat-shock at day 9 of development (Ayroles et al., 2015) |
| darkreared | flies reared in darkness |
| gal80heatshock | flies subjected to heat-shock post eclosion, prior to behavioral assay |
| grownat18 | flies reared in incubators at 18°C |
| grownat20 | flies reared in incubators at 20°C |
| grownat23 | flies reared in incubators at 23°C |
| grownat25 | flies reared in incubators at 25°C |
| grownat30 | flies reared in incubators at 30°C |
| heritability | flies are the progeny of single parents selected for turn biases (Buchanan et al., 2015) |
| intenseenrichment | flies reared in high intensity enrichment population cage (Akhund-Zade et al., 2019) |
| irtest | fly behavior was measured using infrared rather than white illumination |
| mildenrichment | flies reared in mild intensity enrichment vials (Akhund-Zade et al., 2019) |
| normal | standard rearing conditions |
| potato | flies reared on potato media |
| ru486 | flies reared on media supplemented with ru486 |
| ru486control | flies reared on ru486 control media |
| single | flies reared in single housing |
| age | middle of range of ages post-eclosion of fly in that experimental group. E.g., age = 6 typically reflects experimental flies ranging from 4-8 days old |
| experimenterID | name of experimenter who collected the behavioral data |
| trayID | identifying # of the arena array tray in which the fly behaved |
| boxID | identifying # of the imaging box in which the fly behaved |
| acquisition | software used to collect that fly's behavioral data |
| analysis | software used to compute that fly's behavioral measures |
| sex | fly's sex. "both" indicates that both males and females were used in this experimental group, in unspecified proportion |
| eyeColor | state of the white genetic locus. See Fig 5. + indicates wild type, - null, and m mini-white alleles |

**Figure 1.**
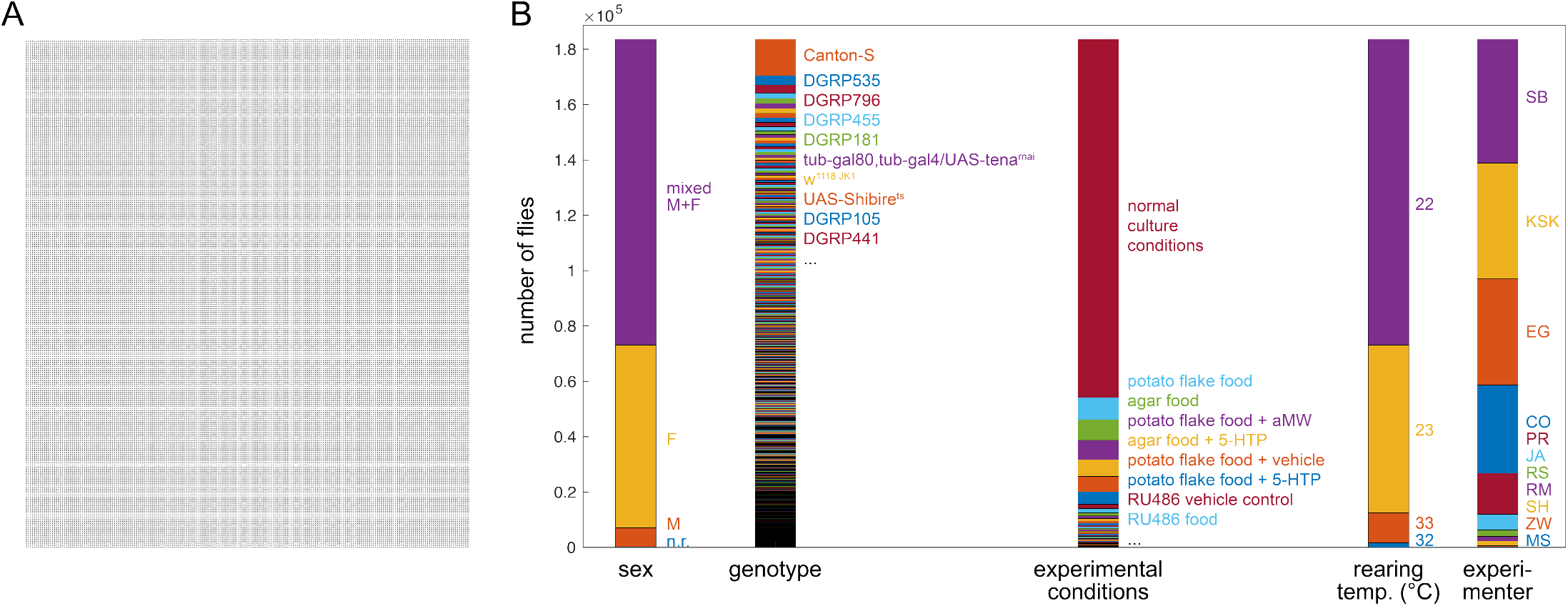
Description of grand Y-maze data set — **A**) Visualization of 183,496 flies (each dot is a fly). **B**) Breakdown of flies into important metadata categories. Height of each color segment indicates the number of flies with that metadata value. Bars align to (A).

The size of our data sets allows some of the most precise estimation of behavioral distributions across individuals to-date. We computed kernel density estimates of the distributions of turn bias, number of turns and turn switchiness (Figure 2A, D, G). The distributions of all measures are essentially unimodal, with the distribution of handedness appearing roughly Gaussian (Figure 2A). However, it deviates from that distribution in a number of ways: it is denser at its mode and in tails corresponding to strong turning biases around 0.1 and 0.9. This is reflected in a kurtosis greater than three (Figure 2B; see below). The empirical distribution of handedness is technically trimodal, with small peaks corresponding to flies with biases very close to 0 and 1. Most flies in these peaks performed fewer than 50 turns, indicating that these peaks may be the consequence of undersampling within these individuals.

**Figure 2.**
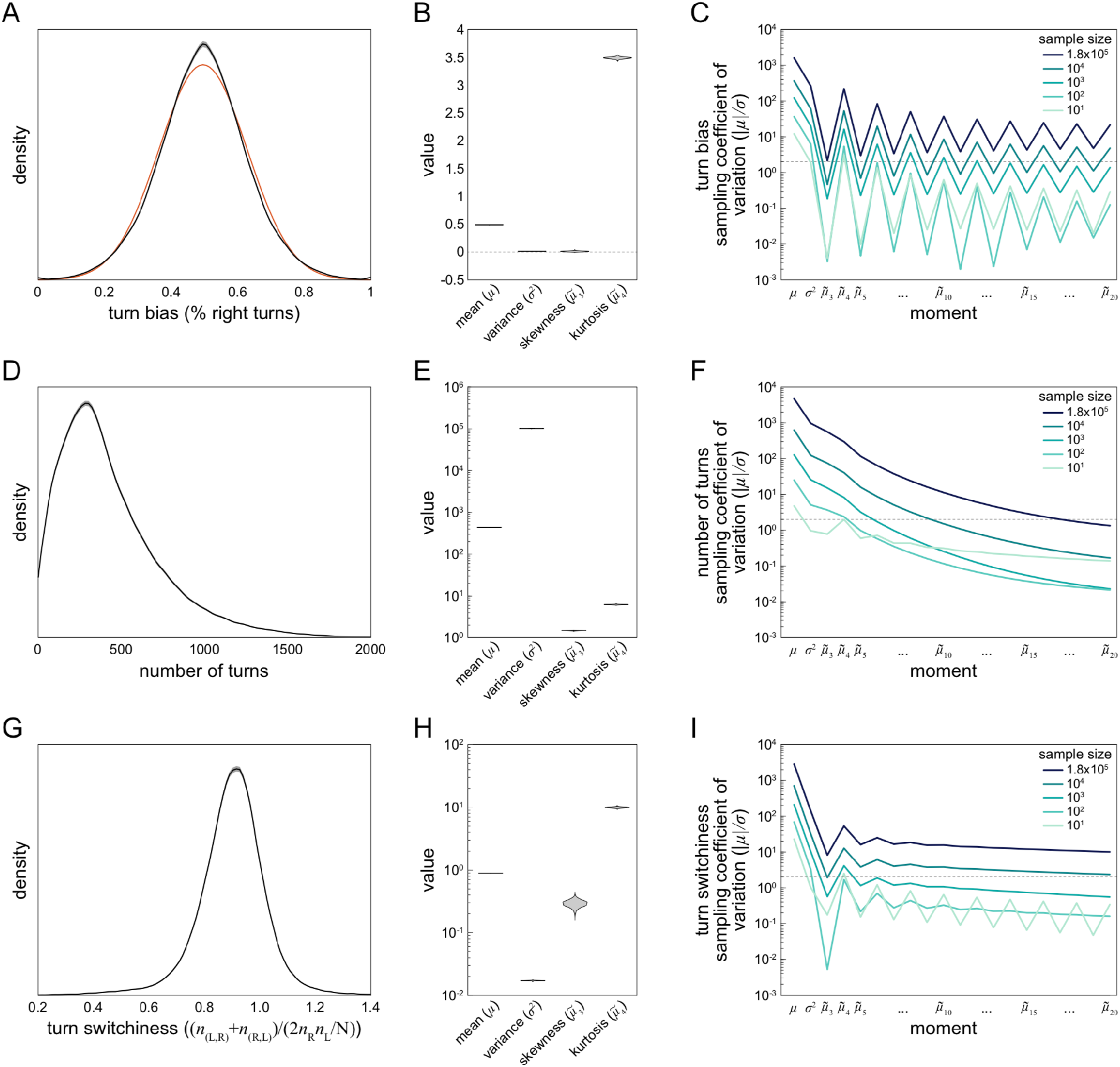
Estimation of statistics describing three Y-maze behavioral measures — **A**) Kernel density estimate of the distribution of turn bias across all flies in the data set. Grey interval is the 95% CI as estimated by bootstrap resampling. Orange line is the Gaussian distribution that best fits the data. **B**) Violin plot of estimation distributions of four statistical moments describing the distribution of turn bias. Each violin is a kernel density estimate of the distribution of each statistic’s value across bootstrap samples from 1000 replicates. **C**) Average bootstrap estimate of the mean, variance and subsequent 18 standardized moments of the distribution of turn bias, as a function of the size of the data set. Darkest line corresponds to the complete grand Y-maze data set, and lighter lines random subsets. Dotted line at |μ|/σ = 2 indicates the threshold for moment estimate significantly different from 0 at p = 0.05. **D-F**) As in (A-C) for number of turns as the behavioral measure. **G-I**) As in (A-C) for turn switchiness as the behavioral measure. Note log y-axes in (E,F,H,I).

To assess the precision of measures quantifying these distributions we looked at the distribution of estimates (under bootstrapping) of the mean, standard deviation, skewness and kurtosis of the behavioral distributions (Figure 2B, E, H). These were generally quite narrow, indicating precise estimation, and generally broader for the higher-order statistics. This was expected as the higher-order statistics have exponential terms that render them more sensitive to sampling error. But their precision did not always decrease monotonically (Figure 2H). To extend this analysis, we computed the standardized moments of each distribution, up to the 20th moment, for each behavioral measure (Figure 2C, F, I). To our surprise, the data provided robust estimates even of the 20th moment of turn bias and turn switchiness. This was true even in 10-fold subsamples of the turn switchiness data, but was not the case for number of turns (Figure 2F) or odd moments of the turn bias data (Figure 2C). This indicates that the reliability of estimates of high-order distribution statistics depends on the underlying distribution, not just the sample size.

In our studies of turn bias in Y-mazes (Buchanan et al., 2015; Ayroles et al., 2015; Akhund-Zade et al., 2019; Werkhoven et al., 2021), we operated under the assumption that the mean turn bias was 0.5 in all genotypes. For example, this assumption was the basis of the decision to not model the interaction of genetic variation for the mean and variability of turn bias in Ayroles et al. 2015. On close examination of this measure in our new data set, we found evidence that the mean turn bias may not be 0.5 (Figure 3). The mean of turn bias in the grand data set was 0.496 (Figure 3A), indicating a slight left bias to Y-maze turn choices. This slight left bias was also present in the distribution of genotype, sex and experimenter (Figure 3B-D) mean turn biases, suggesting that the apparent left bias in the grand mean is not likely attributable to imbalance among the metadata covariates. Indeed, a linear model with 11 meta variables as predictors (all but date, which renders the model rank deficient) and 636 coefficients has a turn bias intercept of 0.485 (SE 0.0099). Moreover, 47/569 genotypes have significant effects in a linear model where genotype is the sole predictor of turn bias (Figure 3E). This is a significant enrichment, and supports the conclusion that the average turn bias is under biological control.

**Figure 3.**
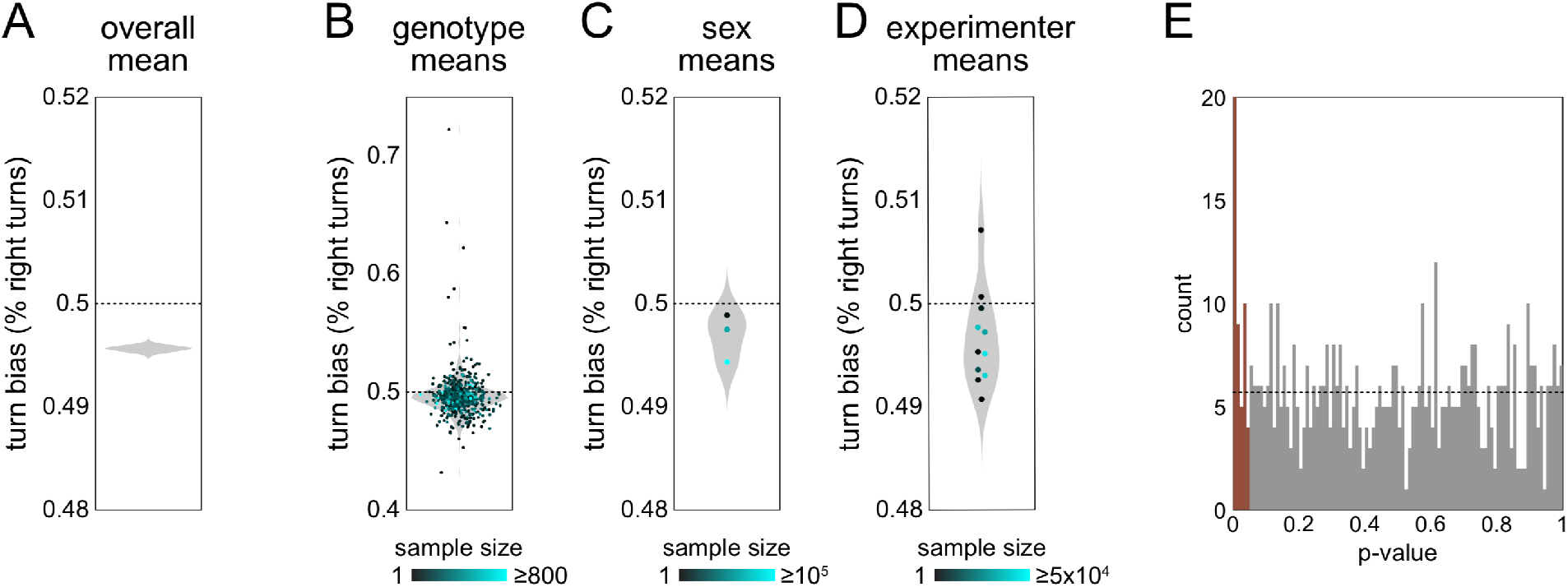
Mean turn bias appears slightly asymmetrical — **A**) Violin plot of estimate distribution for the mean of turn bias across the grand data set, exhibiting an apparent slight left-bias of 49.6%. Violin is a kernel density estimate (KDE) of this statistic from 1000 bootstrap replicates. **B**) Mean turn bias for each genotype (points). Violin is the KDE of genotype means. Point color value indicates the number of flies recorded for that genotype. **C**) As in (B), but with flies grouped by sex experimental condition. The three points correspond, from top to bottom, to males only, females only and mixed sex. **D**) As in (B), but with flies grouped by experimenter. **E**) Histogram of p-values from a linear model with each genotype as a predictor. Brown bars represent effects significant at p < 0.05. Dotted line indicates the expected distribution under the null model.

Since our behavioral data was multidimensional (turn bias, number of turns and turn switchiness were measured for each fly), we were also able to investigate the joint distributions and correlations of these measures. We first tested whether there might be a correlation between turn bias and number of turns, specifically a negative correlation arising from higher sampling error in estimating turn bias for flies making fewer turns. Counter to this prediction, we observed a slight positive correlation (r = 0.036; p = 4*10^−52^). We did, however, notice the effects of the discreteness of number of turns as a measure, and the resulting limited values that turn bias can take on, as a fractal-like (Trifinov et al., 2011) structure in the scatter plot of absolute turn bias vs. number of turns (Figure 4A).

**Figure 4.**
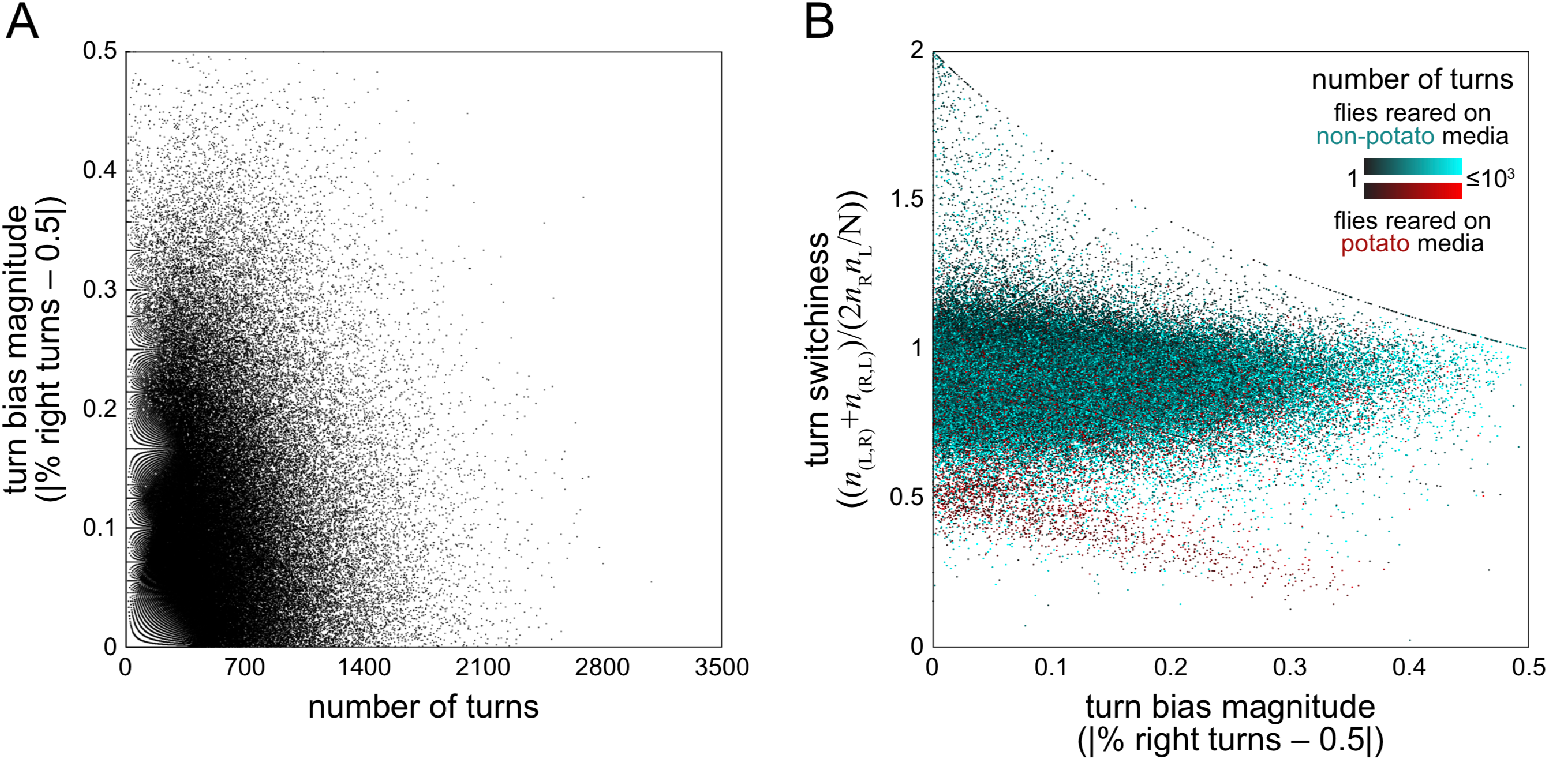
Correlations between behavior measures — **A**) Turn bias magnitude vs. number of turns. Each point is a fly. Fractal-like pattern at left is a consequence of the limited turn bias values that are possible for a given discrete number of turns. r = 0.0357, p < 10-50. **B**) Turn switchiness vs. turn bias magnitude. Each point is a fly and colored on a scale depending on whether the flies were reared on cornmeal-dextrose agar media (black-cyan) or F4-24 potato flake media (black-cyan). Point color value indicates sample size, with dark flies making fewer turns. Curvilinear features are a consequence of limited switchiness values possible for a given turn bias magnitude, a constraint that arises most obviously in flies making fewer turns (dark points).

Next, we examined the joint distribution of turn switchiness and number of turns (Figure 4B). This 2-dimensional distribution had two conspicuous features: an uncorrelated mode containing the vast majority of the flies, and a smaller mode exhibiting a negative linear relationship between turn switchiness and number of turns. The flies in this second mode were predominantly reared on potato flake media (which was sometimes supplemented with drugs targeting the neurotransmitter serotonin; Dierick & Greenspan, 2007; Kain et al., 2012; Krams et al., 2021). Notably, being reared on potato food was not a guarantee that a fly fell in this part of the distribution; the vast majority of flies in such rearing conditions fell in the predominant uncorrelated mode of the joint distribution along with flies fed on standard cornmeal-dextrose media.

Finally, we used the Y-maze data set to revisit several previously examined hypotheses about the proximate mechanisms regulating behavioral variability. We first asked whether the distribution of measures of turn bias variability across genotypes was consistent between the distribution seen in Ayroles et al., 2015 and the other genotypes present in our data set. The lines examined in that paper come from the Drosophila Genome Reference Panel (DGRP; Mackay et al., 2012), a collection of inbred lines established from the natural population of flies in Raleigh, NC USA. The remaining 339 genotypes in our data set come from a variety of sources, mostly lab stocks, and include 165 lines expressing the transgenic driver Gal4 (Brand & Perrimon, 1988) in neural circuit elements (Jennett et al., 2012). Thus, these genotypes do not represent a sample from a natural population. The distribution of their genotype-wise variability in turn bias was largely similar to that observed in DGRP lines (Figure 5A), with genotypes exhibiting coefficients of variation in handedness ranging from less than 0.2 to more than 0.4.

**Figure 5.**
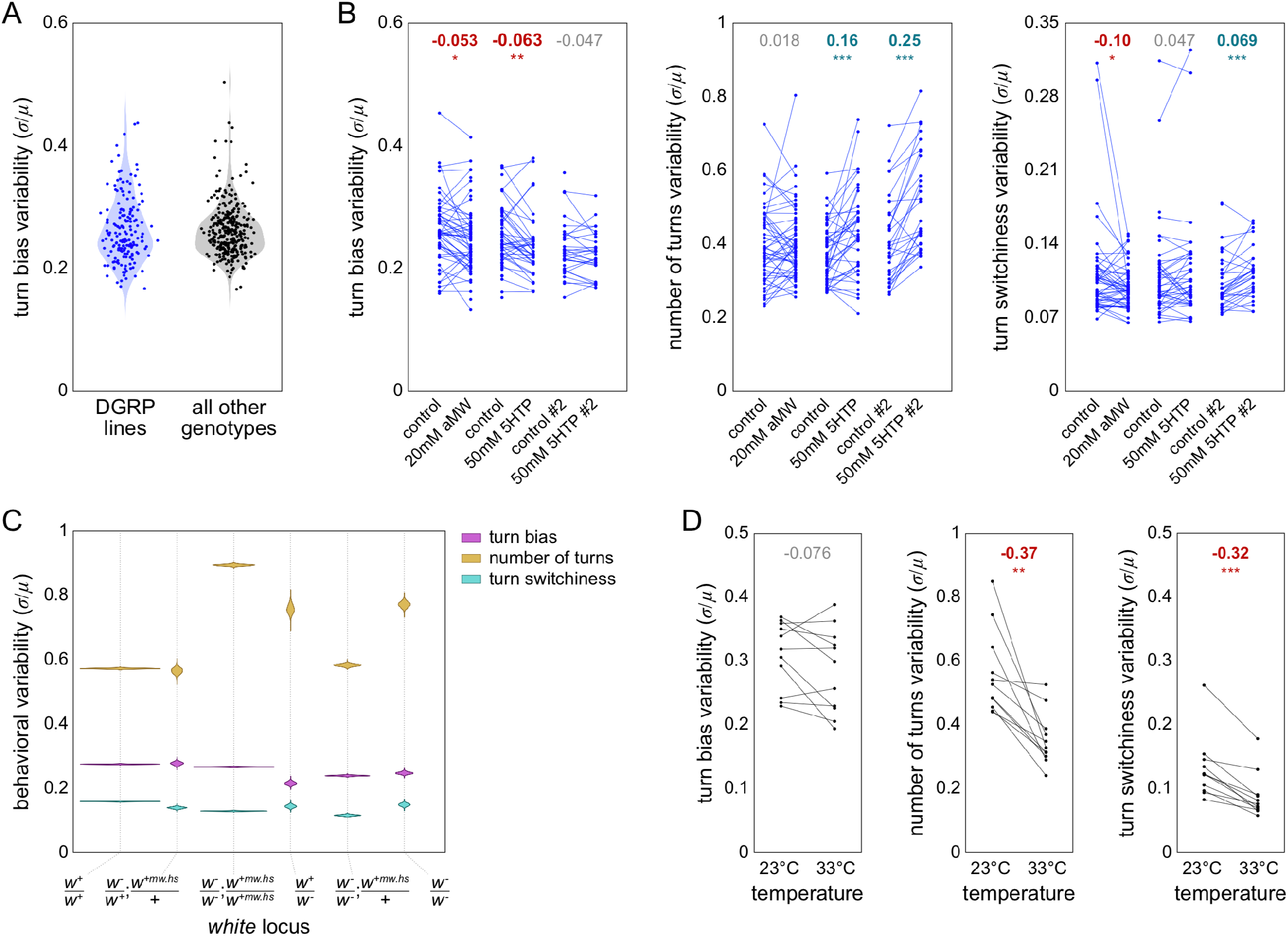
Factors potentially affecting behavioral variability — **A**) Variability (measured as the coefficient of variation) of turn bias for DGRP genotypes (blue) and non-DGRP genotypes (black). Violins are the KDE of genotype variabilities. **B**) Variability of turn bias (left), number of turns (middle) and turn switchiness (right) for DGRP genotypes in 6 pharmacological experimental conditions targeting serotonin. Each point is a genotype in a particular experimental condition. Lines pair genotypes across a drug medium and its associated control medium. Numbers at top indicate the effect size from control to drug treatment. Bold effect sizes are statistically significant and colored by the direction of their effect (red = lower variability; cyan = higher). *: p < 0.05; **: p < 0.01; ***: p < 0.001. **C**) Violin plot of estimation distributions for the variability of turn bias (magenta), number of turns (gold) and turn switchiness (turquoise) vs. genotype of the white gene. *+* indicates wild type, *+mw.hs* the “mini-white” allele typically used to mark a transgenic insertion, and - a null allele (typically *w^1118^*; Hazelrigg et al., 1984). *white* genotypes are roughly ranked in order of expression disruption. The site of *w+mw.hs* insertion varied by line; the semi-colon notation in the panel label indicates that this site might be on a different chromosome than the endogenous *w* locus. **D**) Variability of turn bias (left), number of turns (middle) and turn switchiness (right) for genotypes tested at 23°C and 33°C. Lines pair genotypes across temperature conditions. Effect sizes and significances as in (B).

Neuromodulation may have a special role in the control of behavioral variability (Maloney 2021), e.g., phototaxis (Kain et al., 2012; Krams et al., 2021) and olfactory preference (Honegger & Smith, 2019). We conducted experiments to see if serotonin modulation controls variability of locomotor behaviors in the Y-maze. Specifically, we measured the variability of turn bias, number of turns and turn switchiness in DGRP lines which were treated with alpha-MW (a serotonin synthesis inhibitor), 5-HTP (a biosynthetic precursor of serotonin) (Dierick & Greenspan, 2007) or their respective control media. These treatments generally had small effects on behavioral variability across genotypes (ranging from a −10% to a 7% increase), with the exception of the effect of 5-HTP on variability in the number turns, which, in two versions of the experiment increased variability by 16% or 25% (Figure 5B). Overall, these results imply that although serotonin levels can affect the variability of turn number, there is not a strong effect that is consistent across behaviors.

We previously determined that the effect of serotonin on phototactic variability was dependent on the gene *white*, which encodes a transmembrane transporter that imports the serotonin precursor tryptophan into neurons. We scored the flies in our Y-maze data set for their white genotype, which could range from wild type to homozygous null, with intermediate conditions of (likely) partial rescue by the expression of the “mini-white” allele at non-endogenous transgenic insertion sites (Klemenz et al., 1987). Lines with homozygous null alleles at the endogenous white locus exhibited higher variability in number of turns, with the exception of lines that were also heterozygous for mini-white at a transgenic locus. The molecular function of White suggests that its disruption should produce a behavioral phenotype like serotonin synthesis inhibition, which had no effect in our pharmacological manipulations (whereas feeding flies serotonin precursor increased variability, like *white* disruption). *white* genetic disruption was associated with small reductions in variability in turn bias and turn switchiness (Figure 5C), consistent with the small decreases seen in the aMW pharmacological experiments (Figure 4B). Overall, we found some agreement in the effects of serotonin pharmacological experiments and *white* disruption, but not perfect agreement, suggestive of behavior-dependent complexity in the relationship between *white*, serotonin, and variability.

It has been hypothesized that individuals of the heterogametic sex will exhibit greater trait variability due to noise in gene compensation (James, 1973), though a recent meta analysis found no significant sex-bias in the variances of 218 mouse traits (Zajitschek et al., 2020). We fit linear models to Levene-transformed turn bias, number of turns, and turn switchiness data with genotype and sex as predictors to test for the effect of sex on behavioral variability. Males had variability that was −6.8% (p<0.001), 7.5% (p<0.001) and 1.8% (n.s.) greater than that of females in turn bias, number of turns, and turn switchiness respectively.

Lastly, we examined the effect of temperature during behavioral testing, with the hypothesis that flies would exhibit higher variability at high temperature (32-33°C) than at room temperature (22-23°C). This would be consistent with a mechanism in which heat pushes neural circuits out of the range in which physiologi^−^ cal buffering keeps circuits operating similarly despite latent developmental and genetic variability (Tang et al., 2012; Rinberg et al., 2013). We examined this specifically for genotypes that had paired experiments at low and high temperature, and did not express any temperature-sensitive effectors. We found that high temperature had no effect on turn bias variability, but significantly decreased number of turns variability and turn switchiness variability by 37% and 32% respectively (Figure 5D). Temperature does affect the mean number of turns, typically increasing it by making flies more active. Our analysis controlled for this by assessing mean-normalized variability (the coefficient of variation: μ/σ). Overall, our analyses of the effects of potential proximate mechanisms controlling variability revealed a complex picture with (often small) effects of serotonergic regulation, white genotype, sex and temperature. For all of these manipulations, the direction of effect on variability was behavior-dependent.

## Discussion

We collected Y-maze data collected by lab members back to the origination of this assay 11 years ago. This large data set comprised the behavioral measures of over 180,000 individual flies that made a total of nearly 80 million left-right choices. With it, we were able to estimate the distribution of three measures of individual behavior with unprecedented precision, even out to the 20th standardized statistical moment (Figure 2). In exploratory analyses, we noticed two surprising patterns: 1) a discrete change in the relationship between turn bias magnitude and turn switchiness in a subset of animals that had been reared on potato flake media used for pharmacological experiments, and 2) that flies appear to have a slight left bias in their Y-maze choices. Finally, we used our data set to test several hypotheses pertaining to proximate control of variability in behavior, finding significant behavior-dependent effects of drugs targeting serotonin, mutation of the *white* gene (which encodes a channel that imports serotonin precursor), sex and temperature. Compared to the effects of genotype and the choice of behavior measure, the effects of these manipulations were generally small and context-dependent, underscoring the complexity of relationships between axes of biological regulation and behavioral variability.

Admittedly, a motivation for this study was the desire to explore a very large data set reflecting the work over many years of many lab colleagues. In that spirit, it is fun to think about how throughput might be expanded another order of magnitude in the coming years. One possibility is robotic fly-handling (Alisch et al., 2018), which has yet to be deployed at scale in support of a large screen. Another possibility is tracking flies using capacitive sensors (Itskov et al., 2014) instead of with cameras. This would remove the need for long optical axes that force our behavior boxes to be tall, allowing a dense, vertical packing of arenas within a minimal bench footprint.

While increasing throughput through further automation is an appealing possibility, and perhaps essential for certain classes of experiments (like experimental selection for variability, which would require testing thousands of individual flies per generation for a year or more), it is not without conceptual consequences. One of these is how to assess small effects that are extremely statistically significant due to large sample sizes. Two examples from this study are the apparent slight left turn bias (Figure 3) and the significant positive correlation between turn bias magnitude and number of turns (FIgure 4A). A turn bias of 0.496 compared to an expected value of 0.5 is indeed a small discrepancy, but it might nevertheless be biologically significant given the consistent failure of artificial selection experiments to evolve directional asymmetry in a variety of fly morphological characters (Carter et al., 2009). Another aspect of working with large data sets is that sampling error is likely to be small compared to inadvertent biases in the data (Meng 2018; see Bradley et al., 2021 for an important example). I.e., accuracy is unlikely to improve with further observations, but instead with the harder work of identifying systematic miscalibration, misunderstandings of what is being measured, or structure in the data leading to effects like Simpson’s paradox. A way forward among these challenges may be to conduct experiments and analyses under a variety of biological conditions, increasing the odds that inferences generalize across contexts (Voelkl et al., 2020), an approach that would also be boosted by throughput and automation.

With caveats of big data in mind, we want to consider possible errors that might explain the apparent slight (but highly significant) left mean turn bias. All experimenters conducting these experiments are right-handed. It is formally possible that chiral manipulation during the experimental set-up imparted a slight chirality to turning in the Y-maze, though we cannot think of a convincing mechanism by which this would happen. We can also not think of mechanisms by which small, inevitable asymmetries in our behavioral rigs would impart a consistent left bias to behaviors measured across several generations of rigs and tracking software versions. Arguments in favor of the apparent left turn bias being real are previous reports of small mean asymmetries in wing size and shape (Klingenberg et al., 1998), possible indirect effects of conspicuously asymmetrical anatomical features like the gut, or the contribution of the Asymmetric Body, a small neuropil abutting the premotor Central Complex that is consistently larger in the right hemisphere (Wolff & Rubin, 2018).

While we found that our data set allowed the precise estimation of the distribution of individual behavioral scores, we also saw that the stability of higher-order moment estimates depended strongly on the behavioral distribution in question (Figure 2). Thus, there is not necessarily a simple rule for how large a sample should be to estimate higher order statistics of its distribution. In the joint distribution of turn bias magnitude and turn switchiness, we observed two distinct modes between these measures, and, to our surprise, found that most of the points falling in the rarer mode came from experiments where flies were reared on potato flake food (Figure 4B). These flies comprised a relatively small subset of multiple experiments, in both control and drug conditions, from many genotypes. Thus, rearing on potato media is the best explanatory variable we could find for this mode of variation. We previously observed that acutely switching flies from cornmeal-dextrose media to potato media increased their variability in odor preference (Honegger & Smith, 2018). Perhaps this perturbation also alters the correlation structure, in a subset of flies, between turn bias and turn switchiness. Since these measures may relate to the paths animals take through natural environments, a food-dependent change in turning might alter foraging statistics, perhaps adaptively.

Finally, we used this large data set to examine hypotheses about proximate mechanisms controlling variability. We found many significant effects, such as 5-HTP or disruption of the *white* locus increasing variability in number of turns, disruption of white decreasing variability of turn bias and turn switchiness, males exhibiting slightly lower variability in turn bias but higher variability in number of turns, and conducting experiments at high temperatures lowering variability in number of turns and turn switchiness (Figure 5). We expected temperature to increase variability per results in the crab stomatogastric ganglion (Tang et al., 2012; Rinberg et al., 2013), but our high temperature experiments did not push the flies to their critical thermal limits (Kellermann et al., 2012). Thus, perhaps even higher temperature manipulations might result in consistent increases in variability across behaviors. Our variability results indicate a complex, behavior-dependent relationship between many biological mechanisms and behavioral variability, which likely parallels the complexity of mechanisms controlling the means of behavioral traits. Experimental automation, and the high throughput it permits, made these and other findings on behavioral individuality feasible.

## Methods

### Data and analysis code

All behavioral measures and metadata values, along with the code underlying analyses are available at *http://lab.debivort.org/precise-quantification-of-behavioral-individuality*/ and *https://zenodo.org/record/5784716*.

### Fly handling

Unless otherwise indicated, the default culture conditions were cornmeal-dextrose media and incubation on the bench or in incubators at 21°C-25°C with 12h/12h light cycles. Flies were generally anesthetized under CO_2_ to load them into y-mazes, though a small portion of flies were anesthetized by ice or loaded without anesthetization. Flies were given a period of 15-30 minutes of acclimation to the Y-mazes after loading before data collection began.

### Pharmacological experiments

Experimental flies receiving drug treatments were reared from egg-laying in drug-supplemented media (or control media). Drug media are indicated in the expCond metadata variable (see Table 1). To supplement media, drug was added to water, which was then added to dry potato flake media, or drug was added to agar media liquified momentarily in a microwave oven. 15mg ascorbic acid was added to each 60mL media vial as an anti-oxidant in 5-HTP treated groups and their controls. The two 5-HTP experiments presented in Figure 5 were conducted on potato media and cornmeal-dextrose media (#2) but are otherwise identical. To control for the average dose of experimental flies, prior to drug experiments we measured the average number of progeny to eclose following a 24h parental egg-laying session, on cornmeal-dextrose media, for each of the DGRP lines. The number of parental animals for drug experiments was adjusted proportionally, line-by-line, to target an identical number of progeny on the drug media for each line.

### Behavioral assays

Data was collected in Y-shaped mazes arrayed in trays (Buchanan et al., 2015; Alisch et al., 2018; Werkhoven et al., 2019) and imaged in enclosed behavioral boxes (Werkhoven et al., 2019) under diffuse white LED illumination typically provided by custom LED boards (Knema LLC, Shreveport, LA USA). The default assay length was 2h. Fly centroids were computed in real time using background subtraction implemented in a variety of custom software environments coded in LabView or MATLAB. The centroid tracking software used in recent experiments was MARGO (Werkhoven et al., 2019).

### Statistics and analysis

Analysis was conducted in MATLAB 2017b (The Mathworks, Natick, MA USA) using custom functions. 95% confidence intervals estimated by bootstrapping were estimated as +/− twice the standard deviation of values across bootstrap replicates. For the analysis of the effect of temperature on variability (Figure 5D), the 23°C groups include experiments conducted at 22°C and the 33°C groups include experiments conducted at 32°C. Genotypes were only included in the temperature analysis if they had data recorded at both temperatures and did not express any thermogenetic constructs. Thus, most genotypes in this analysis were controls of thermogenetic experiments or wild type lines. Significance in the serotonin pharmacological and temperature experiments was assessed by paired t-tests, and all reported p-values are nominal.

## Acknowledgements

We thank Ed Soucy, Joel Greenwood, and Brett Graham of Harvard’s Neurotechnology Core for their help with instrument engineering. BdB was supported by a Sloan Research Fellowship, a Klingenstein-Simons Fellowship Award, a Smith Family Odyssey Award, a Harvard/MIT Basic Neuroscience Grant, National Science Foundation grant no. IOS-1557913, and NIH/ NINDS grant no. 1R01NS121874-01. KSK and ZW were supported by NSF Graduate Research Fellowships #DGE2013170544 and #DGE1144152. JAZ and MAYS were supported by the Harvard Quantitative Biology Initiative. JAZ was supported by The NSF-Simons Center for Mathematical and Statistical Analysis of Biology at Harvard, award number #1764269.

